# How does temperature affect the dynamics of SARS-CoV-2 M proteins? Insights from Molecular Dynamics Simulations

**DOI:** 10.1101/2021.10.05.463008

**Authors:** Soumya Lipsa Rath, Madhusmita Tripathy, Nabanita Mandal

## Abstract

Enveloped viruses, in general, have several transmembrane proteins and glycoproteins, which assist the virus in entry and attachment onto the host cells. These proteins also play a significant role in determining the shape and size of the newly formed virus particles. The lipid membrane and the embedded proteins affect each other in non-trivial ways during the course of the viral life cycle. Unravelling the nature of the protein-protein and protein-lipid interactions, under various environmental and physiological conditions, could therefore prove to be crucial in development of therapeutics. Here, we study the M protein of SARS-CoV-2 to understand the effect of temperature on the properties of the protein-membrane system. The membrane embedded dimeric M proteins were studied using atomistic and coarse-grained molecular dynamics simulations at temperatures ranging between 10 and 50 °C. While temperature induced fluctuations should be monotonic, we observe a steady rise in the protein dynamics up to 40 °C, beyond which it surprisingly reverts back to the low temperature behaviour. Detailed investigation reveals disordering of the membrane lipids in the presence of the protein, which induces additional curvature around the transmembrane region. Coarse-grained simulations indicate temperature dependent aggregation of M protein dimers. Our study clearly indicates that the dynamics of membrane lipids and integral M protein of SARS-CoV-2 enables it to better associate and aggregate only at a certain temperature range (i.e., ~30 to 40 °C). This can have important implications in the protein aggregation and subsequent viral budding/fission processes.

## Introduction

A deadlier form of severe acute respiratory syndrome (SARS) virus, reappeared in 2019 and was named as SARS-CoV-2 [1]. The virus is highly contagious and detrimental to the respiratory tissues and ancillary organs of humans [2]. The Covid19 pandemic caused by SARS-CoV-2, has so far resulted in a large number of fatalities and numerous casualties are still being reported around the world [1, 2]. Similar to other coronaviruses, it encodes four major structural proteins; spike (S), membrane (M), envelope (E) and nucleocapsid (N). The S protein is the main interaction site for the human receptor, Angiotensin Converting Enzyme 2 (or ACE-2), the N protein associates with the RNA present in the viral genome, the M and E proteins are found on the lipid envelope and help in assembly and release [2, 3].

The viral M protein is one of the most abundant structural proteins distributed on the surface of the viral envelope. It associates with the S, E and N proteins and directs the virus assembly and aids in the budding process [4]. During synthesis, S, E and N proteins are translated and inserted into the Endoplasmic Reticulum (ER) of the host cell, from where it is transferred to the Golgi complexes with the help of the M protein [5, 6, 7, 8, 9]. The E and M proteins trigger the virion budding, where the nucleocapsid is already present in an enclosed form. The S proteins are subsequently incorporated on these virions. These large vesicles are then released from the host cells through exocytosis. Thus, the M proteins play a crucial role in the viral assembly and release [10, 11, 12]. The co-expression of M and E proteins alone is sufficient to trigger the formation of virions as observed in insect cells [13, 14, 15]. However, the majority of the research on SARS virus has so far focused exclusively on the S proteins [16]. While the extensive research has led to successful development of a number of effective vaccines, one of the fastest in medical history, our knowledge on the role and importance of other structural proteins and their interaction still remains inadequate. Only recently, there have been some noticeable attempts along this line: both towards modeling the other integral protein structures and assessing their validity in molecular simulation [17,18,19,20]. In this work, we focus our attention on the integral M protein. We investigate the nature of protein-protein and protein-membrane interactions and investigate the effect of temperature on them.

An intriguing aspect of the viral particle formation process is the ability of the virus to exhibit pleomorphism. Pleomorphism is an example of extreme natural variations that allow changes in size and shape of enveloped Viruses such as influenza A virus or arenavirus [21, 22, 23, 24]. The coronaviruses are not an exception to this phenomenon [24, 25]. However, the tolerance to pleomorphism depends on the structural proteins present on the viral envelope. Extreme conditions might result in failed assembly of virions or might lead to the formation of non-infectious viral particles. Although such variations have been reported in literature, very few studies have been carried out to understand how the overall virus structure is shaped [21–25]. Another important factor in the viral particle assembly is the number of M proteins in the membrane. It has been observed that a greater number of M proteins generate larger viral particles [5, 6, 26].

The coronavirus family M proteins share a high level of structural and functional similarity. The structure of M protein comprises a membrane bound domain, an amino terminal, which is the ectodomain, and a carboxy terminal domain, which is towards the nucleocapsid or endodomain [27]. The M proteins largely accumulate in the Golgi apparatus or the ER [11]. Apart from associating with the N, S and E proteins, the M protein is capable of associating with itself forming oligomers. For the development of a functional viral particle the M protein should be able to successfully associate with the N and E proteins [26, 27]. Hence, knowledge of the dynamics and behaviour of the M protein is crucial in the understanding of the formation of the virus particle.

A distinct feature of the coronavirus envelope is its anionic nature [28, 29, 30]. The membrane in enveloped viruses enables them to sustain high environmental temperatures compared to that of non-enveloped viruses [31]. Reports by Polozov et al. show the existence of gel-phase of lipids at temperatures below 22°C, which is independent of the protein content in the membrane. This rubbery, gel-like state of the influenza lipid molecules on the envelope helps them withstand low temperatures and also helps in the airborne transmission of the virus [32]. Although the basic characteristic of the phospholipid bilayer can be independent of the protein that is embedded in the viral lipid envelope, it is interesting to study how the lipid bilayer membrane and M protein together influence the size and shape of the virion particles at different temperatures.

The global pandemic caused by SARS-CoV-2 has been observed at varied temperature conditions, ranging from mere 8-10 °C in China to high ~40 °C in sub-tropical countries like India [33,34]. Earlier studies have shown the influence of temperature on the structural properties of S proteins [35]. Although extensive studies on drug development, mutation and dynamics of SARS-CoV-2 protein are being carried out globally, it is important to understand how the architecture of the pleomorphic coronavirus, shaped by the M proteins, is affected by variations in temperature [16, 36]. Towards this, we first investigate the properties of SARS-CoV-2 membrane protein embedded in a lipid bilayer using atomistic molecular dynamics (MD) simulations. Subsequently, we employ coarse grained (CG) MD simulation to investigate the aggregation behaviour of proteins on relatively larger membrane-protein systems. Our results suggest that the dynamics of the transmembrane domain of the M protein is mostly governed by the state of the lipid membrane, while the ectodomain exhibits distinct dynamics in the physiological temperature ranges (~30 to 40 °C). However, the presence of M protein does not seem to affect the nature of the model anionic membrane. Results from CG MD simulation suggest that the proteins cluster and form stable aggregates only beyond a certain temperature threshold, an observation that can be attributed to the gel phase of the membrane at lower temperatures. Furthermore, while a single M protein dimer can impart local deformations, clustering of M protein dimers does not seem to generate any large-scale deformation in the membrane, which indicates the possible role of other integral proteins along with M protein in the viral budding/fission process. The rest of the paper is organized as follows. We discuss the details of the atomistic and CG simulations in Section 2. In Section 3, we discuss the important results from the study. We conclude in Section 4 along with a brief discussion on the scope of the work and possible future directions.

## Materials and Methods

### Atomistic simulations of an M-protein dimer in a model anionic membrane

The M protein model was obtained from the structures deposited by Feig’s lab [17]. In their work, the protein structure was predicted using inter-residue distance prediction-based modeling using trRosetta, where out of 10 predicted models, the lowest energy conformation was considered for subsequent studies. Alpha Fold models were also used along with trRosetta models as initial structures for machine learning. This was followed by MD simulation-based refinement. In a recent study, Monje-Galvan and Voth [18] have compared the two models of M protein from Feig’s lab based on their stability and the lipid sorting pattern around them. They found that while both the conformations are stable in water as well as in a complex bilayer, the lipid sorting pattern and the resulting membrane deformation profile around them are markedly different. In this study we have used the open conformation model of the M protein [17, 18]. In our study, a lipid bilayer membrane constituting DPPC and DPPG lipids in the ratio of 7:3, embedding a single M protein dimer, was built using CHARMM-GUI builder [37]. The initial coordinates of the lipids were generated by the membrane builder tool of CHARMM-GUI. Water molecules and neutralizing ions (NaCl) were subsequently added to the system, such that the initial box was 10 nm x 10 nm x 13 nm in the X, Y, and Z directions, respectively. The total number of atoms in the system was 133479, with 210 DPPC and 90 DPPG lipids, and 29273 water molecules. The CHARMM36 force field was used for protein and lipid molecules [38] and TIP3P model for water [39]. Standard procedure was followed for the initial minimization and equilibration, and 0.5 µs long production simulations were performed for subsequent analyses. The simulations were performed at five different temperatures, 10°C, 20°C, 30°C, 40°C and 50°C using GROMACS 5.0 molecular dynamics simulation software [40, 41].

### Coarse Grained simulations of the M-proteins in membrane

Coarse grained (CG) simulations were carried out using Martini 2.2 [42] amino acid, Martini 2.0 [43] lipids and non-polarizable water models [43] and Gromacs 5.0 simulation package [41]. We used the Martini Bilayer builder [44] for generating two CG systems, embedding 8 and 64 dimeric M proteins, respectively in a model bilayer. Subsequently, solvent (CG water) and neutralizing Na^+^ and Cl^-^ ions were used to build bilayer-protein systems similar to the previous all-atom simulations. The box size for the 8-dimer system was 40 nm × 40 nm × 13.5 nm and the total number of CG beads was 180231, including 3423 DPPC and 1467 DPPG lipids, and 112172 water beads. Similarly, for the 64-dimer system, the simulation box size was 85 nm × 85 nm × 13.5 nm and the total number of CG beads was 827816, constituting 14791 DPPC and 6339 DPPG lipids, and 493141 water beads. The important system details are summarized in Table S1.

Both the all-atom and CG simulations were carried out using CHARMM36 force-field [37,42] parameters for proteins, lipids, and ions and were prepared using the standard protocol of CHARMM-GUI for bilayer simulations [45–47]. After the initial minimization over 1 ns, equilibration was carried out in the NVT ensemble by gradually reducing the dihedral restraints. Subsequently, equilibration was carried out in the NPT ensemble. Production runs of 0.5 µs duration, in the NPT ensemble, were carried out in the temperature range between 10 to 50 °C (Table S1). All analyses were carried out using Gromacs analysis programs. Pymol [48] and VMD [49] were used for visualization and generating structures. We use the open-source tool g_lomepro [50] to compute the curvature profile of the lipid membrane, which is integrated as an analysis suit in Gromacs. This tool uses a grid-based algorithm (a modified version of GridMAT-MD [51]) to map the lipid atoms on a bilayer membrane and the embedded proteins to two 2D grids, corresponding to each leaflet in the bilayer. Spatial derivatives at each grid point of the surface are then calculated and used to compute the mean and Gaussian curvatures using geometric methods [52, 53]. Furthermore, spectral filtering is used to capture specific modes in membrane curvature. Here we focus our attention on the mean curvature values. We also compute the Area Per Lipid (APL) and thickness using g_lomepro.

## Results and Discussion

The SARS-CoV2 membrane (M) protein is 222-residue long and is embedded in the lipid bilayer. The N-terminal of the M protein is exposed to the external surface of the viral capsid and is in the form of a β sheet [4, 25]. Previous studies have hinted towards a negatively charged viral lipid bilayer [29]. Therefore, in this work, we have used a 7:3 ratio of DPPC: DPPG lipids to mimic the viral membrane as studied in earlier literature [51, 52]. While DPPC is a neutral lipid, DPPG carries a net negative charge. Similar composition of DPPC and DPPG have been earlier reported in pulmonary surfactants and hence used in this study as a model system [51]. Although it is well known that the gel-to-crystalline transition temperature for both DPPC and DPPG lipids is around 41 °C [53, 54], it is yet to be elucidated whether the presence of M protein in the anionic membrane modulates the relative temperature withstanding capabilities or brings about noticeable biophysical changes.

Individual SARS-CoV-2 Membrane proteins resemble a Greek amphora. Structurally, the M protein is organized into a small N-terminal ectodomain, three membrane spanning transmembrane regions and a large carboxy terminal endodomain (Figure 1) [24]. While the monomer-monomer interactions are reported to take place at the transmembrane region, M protein interactions with the S, E and N proteins generally take place at the carboxy terminal domain. The M proteins are usually present as a dimer linked near the transmembrane region [25, 26].

**Figure 1.**
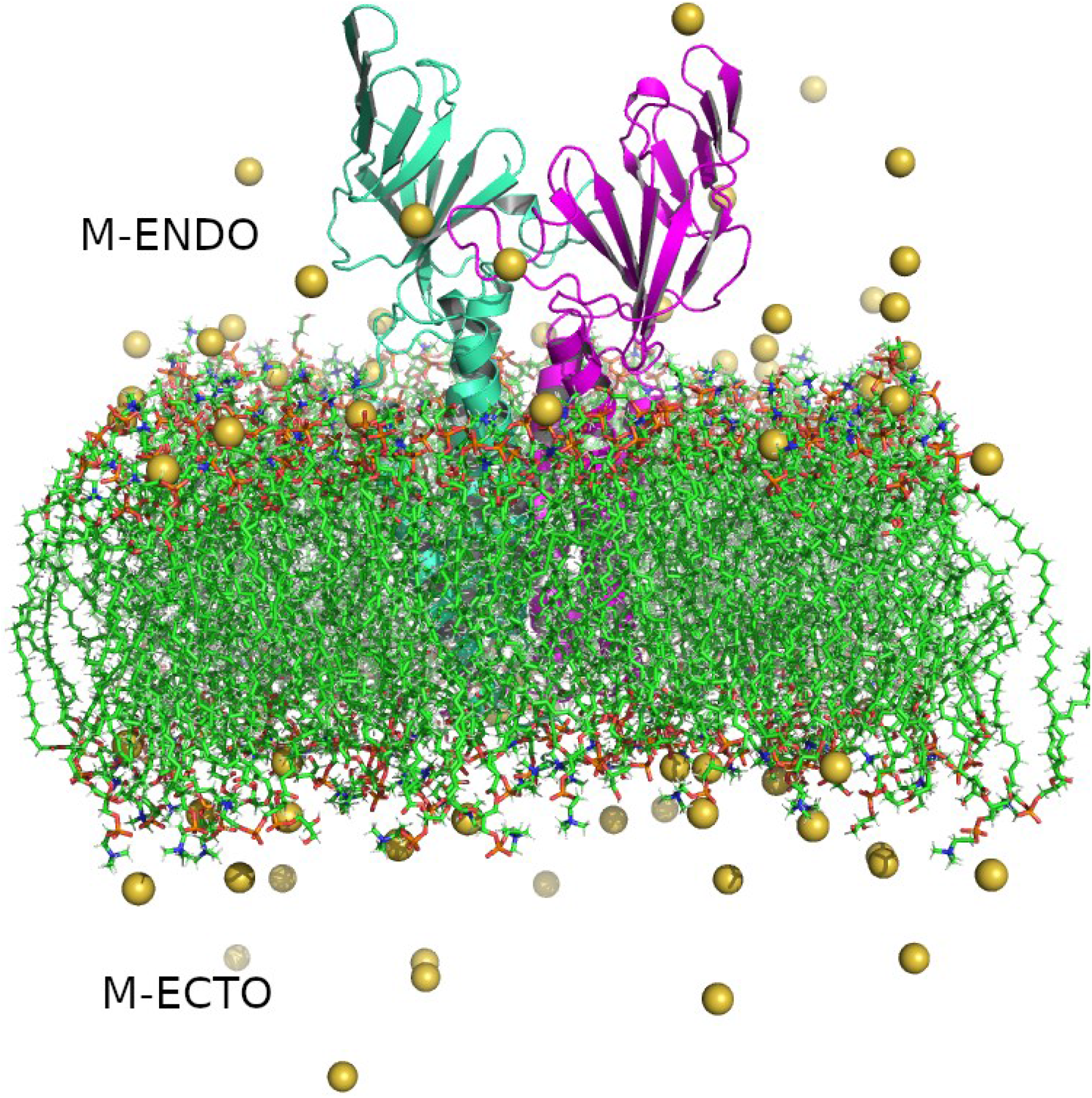
Structure of SARS-CoV-2 Membrane protein dimer. Monomers are shown in cyan and magenta in cartoon representation, lipid bilayers in CPK representation, and neutralizing sodium ions are shown as yellow vdW spheres.

Two forms of the M-protein have been reported [25]. The first form is in an elongated conformation, where the protein is capable of interacting with the nucleoprotein and imparting curvature to the membrane structure: reportedly 5-6 degrees per M protein dimer. The second compact form has very distinct boundaries and generally forms protein aggregates but does not help in the membrane curvature. It is possible that one form can be converted to another and vice versa under suitable environmental conditions [25, 55]. The Feig lab [17] have also reported two structures of the M protein: one based on the trRosetta method and a further refined structure using AlphaFold models. The two structures differ based on the orientation of the extra-membrane N-terminal region with respect to the transmembrane region. In this work, we use the former model of the M protein where the N-terminal domains remain in an open state.

To assess if the system has attained stability, we checked for the Root Mean Square Deviation (RMSD) of the protein at the different temperatures over the simulation time (Figure S1). It was observed that after 150 ns, the systems were largely stable. The RMSDs at 30, 40 and 50 °C were found to be relatively higher than those at 10 and 20 °C. Next, we compared the Root Mean Square Fluctuation (RMSF) of the CA atoms of the protein at various temperatures (Figure 2). The average RMSFs indicate that the protein dynamics remained overall consistent at different temperatures. Distinct peaks were found for residues 9-47 at temperatures 40 and 50 °C. The peak corresponds to the outermost part of the transmembrane helix towards the C-terminal. Comparatively lesser but noticeable changes were also observed for the remaining two helices lying close to the N-terminal. While the overall RMSFs were similar, solvent exposed C-terminal loops around residues 180-190 and 203-220 displayed higher flexibility. Interestingly, the transmembrane helix flexibility is directly correlated to membrane properties such as diffusivity, fluidity, etc [56, 57].

**Figure 2.**
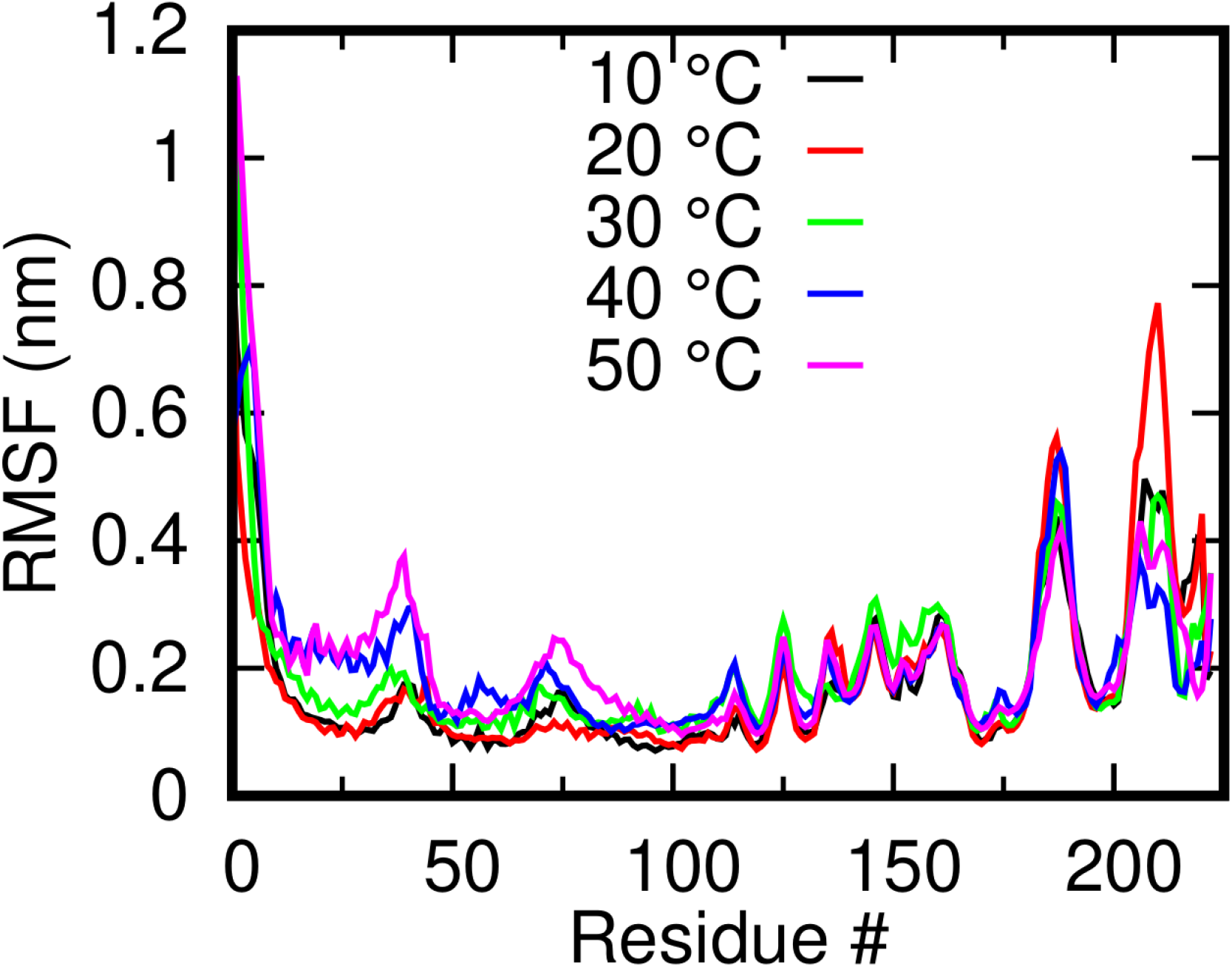
Root Mean Squared Fluctuation of the M-protein monomers (mean of both dimers) at different temperatures shows the overall stability of the systems.

We access the overall temperature effects on the membrane-protein system in terms of 1) the structure and dynamics of a single M protein dimer, 2) the characteristics of the lipid membrane, 3) the ability of the M protein dimer in generating membrane curvature, and 3) the clustering behaviour of multiple M protein dimers.

### Effect of temperature on individual M protein dimers

To investigate the effect of temperature on the structure of the M protein dimer, we calculated the distribution of a pair of distances between the two monomeric units. The CA atoms of residue LYS15 and THR169 were considered as reference sites (see Figure 3). LYS15 lies towards the end of the first helix and stays buried in the lipid membrane, while THR169 lies on the C-terminal β strands and is far from the influence of the membrane. The distance between the two reference sites on each monomeric unit were computed over the production trajectory at every temperature and the 2-dimensional (2D) probability distribution of these two distances were calculated. In Figure 3, we show these 2D distributions at various temperatures, where X-axis represent the distance between LYS15 sites (hereafter, referred as d_15_), Y-axis represent the distance between the THR169 sites (referred as d_169_) of the two monomers, and the colorbar indicate the probability. The ranges of both the axes and the colorbar are kept the same so as to better compare the distributions at various temperatures.

**Figure 3.**
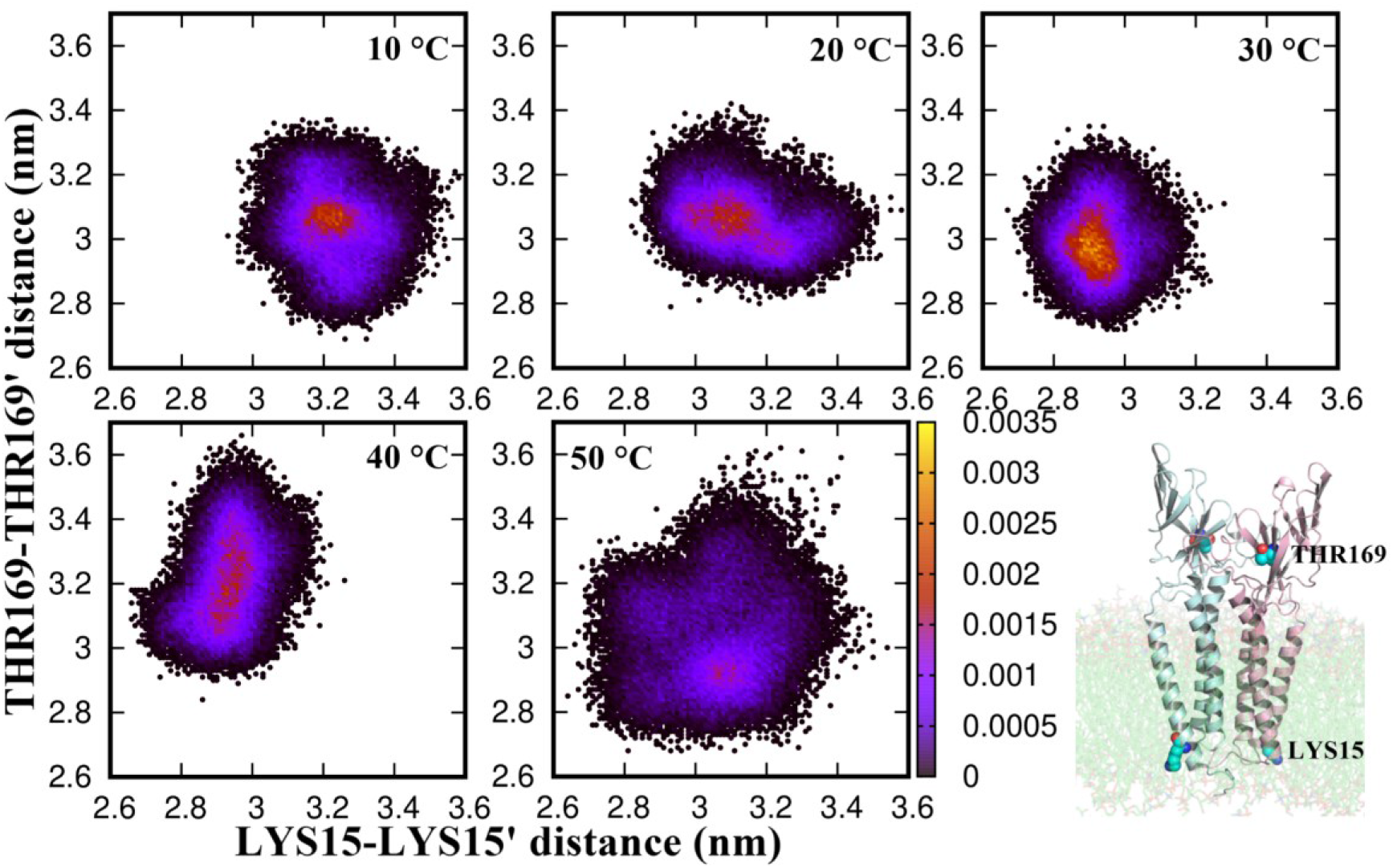
2D distribution of inter-monomer distances of the M protein at different temperatures.

The transmembrane part of the protein dimer is restricted in the gel-like lipid membrane at low temperature. With increasing temperature, the lipids in the membrane become more disordered (discussed below) and the flexibility of the transmembrane domain increases so as to allow it to sample different conformations. As evident, the 2D distribution at 10 °C is almost circular around a central high probability value of 3.2 nm and 3.05 nm along d_15_ and d_169_ axes, respectively. This indicates single narrow peak distributions along the individual axes, which might indicate a meta-stable conformation of the transmembrane domain, rather than a stable one, owing to the low temperature. We observe that d_15_ peaks at <3.1 nm at 20°C and around 2.9 nm at 30 and 40 °C. The distribution along d_169_, which changes very little between 10°C and 30°C (< 1 nm), becomes wide at 40 °C and beyond, indicating large thermal fluctuations of the extrinsic domains. At 50 °C, d_15_ peaks around <3.1 nm (similar to that at 20°C) with a shoulder at 2.9 nm (similar to the peak positions at 30 and 40 °C), and d_169_ peaks around >2.9 nm. However, the distribution is quite broad owing to large thermal fluctuations and the peak is rather shallow as compared to those at other temperatures (see the color bar). The distribution at 50 °C is wide enough to cover all probable distances in d_15_ and d_169_ that are accessible at the lower temperatures. The motion of the protein subunits at high temperature, thus, seems to be governed by the thermal fluctuations rather than indicating any functional motion of the protein. On the other hand, the distribution at 30 °C seems to indicate a stable protein confirmation with sharpest narrow peaks along both axes and a well-defined motion of the N-terminal domains at 40 °C. To further understand the implication of the increased thermal fluctuation in the functional motion of the various protein domains, we subsequently performed the principal component analysis (PCA).

For getting more clarity of the dynamics, the part of the protein embedded in the membrane was not considered for the PC analysis (Figure 4a). It was found that the first few eigenvectors were sufficient to describe the motions of the protein. Figure 4 shows the free energy landscape of the first two principal components using the CA atoms of the extrinsic domain of the M protein. The energy is computed as -k_B_Tln P_(PC1, PC2)_, where P_(PC1,PC2)_ is the distribution probability of the structures at different temperatures. While the protein explored an extended phase space in (PC1, PC2) at 10, 20, and 30 °C, the minimum energy states were rather well defined at 40 and 50 °C. As observed from Figure 4, the ΔG of the most populated conformation at 10, 20, 30, 40 and 50 °C were found to be 18.1, 16.6, 15.2, 20.2 and 19.1 kJ/mol, respectively. At 30 °C, along with a wide and shallow distribution of the conformations, the ΔG value was found to be the lowest. In contrast, a narrow distribution with a high ΔG value was seen for the protein at 40 °C, thus indicating a sharp transition from 30 to 40 °C. From the PCA plots it is clear that at 30 °C the extrinsic domain of protein is very dynamic. On the other hand, a stable conformation was observed at 40 °C that was different from the ones at 10, 20 and 50 °C, clearly indicating the difference in protein dynamics over the physiological temperature range. However, at higher temperatures, although the protein doesn’t undergo any secondary structure change, the increased thermal motion might influence the dynamics of the extrinsic domain of the protein and correspondingly the free energy basins.

**Figure 4.**
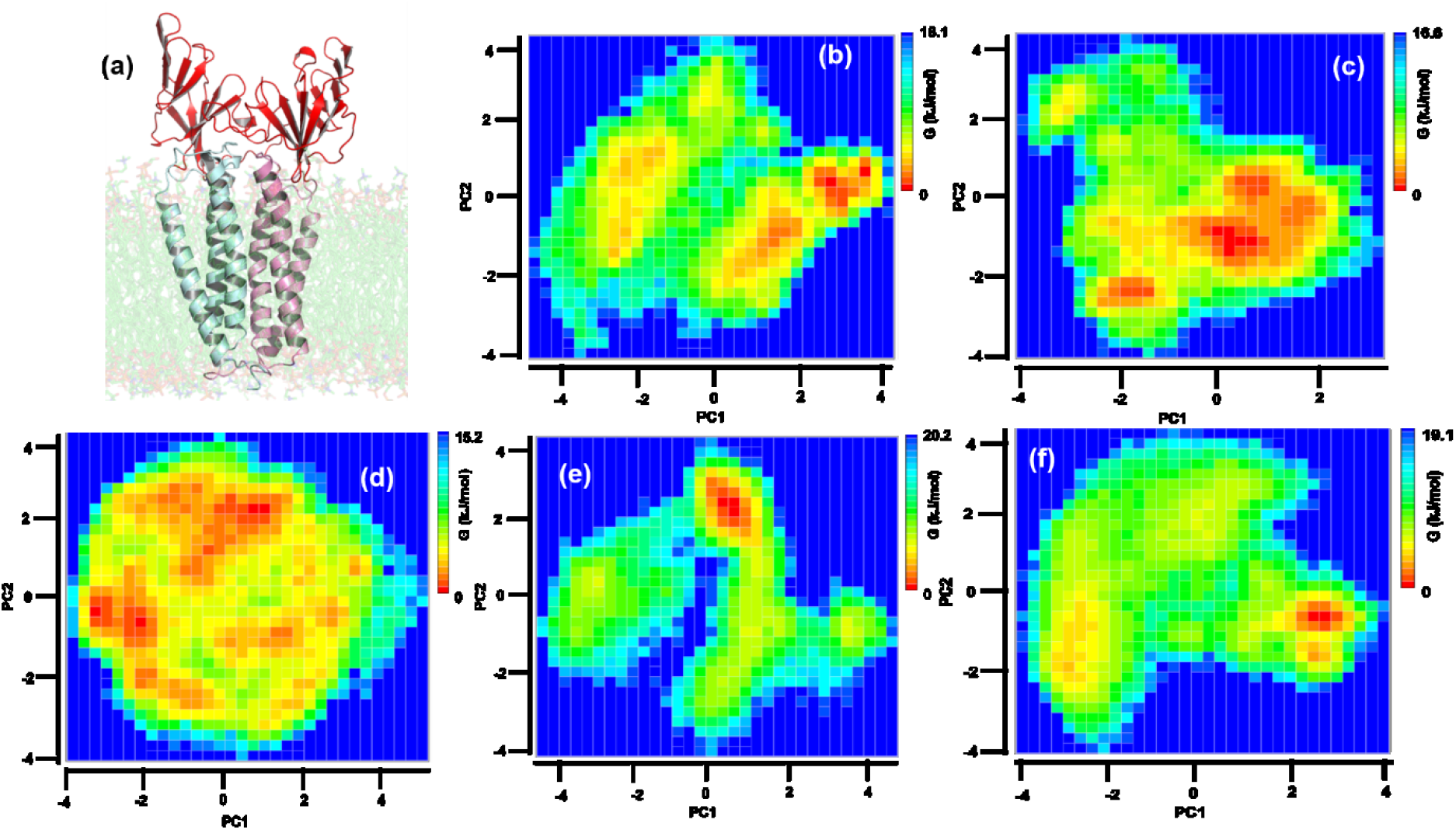
Free Energy landscape of (a) extrinsic domain of M protein (highlighted in red cartoon) at (b) 10 °C, (c) 20 °C, (d) 30 °C, (e) 40 °C and (f) 50 °C. The red color depicts the most stable conformation of the protein.

### Effect of temperature on the embedding lipid membrane

Next, we investigated whether the lipid membrane, in the presence of the M protein, shows patterns similar to earlier observed anionic lipid membranes or does it have the inherent capacity to withstand high temperatures in the presence of protein as reported in the case of Influenza and other related viruses [32]. To monitor this, we computed for the lipid tail order parameter, also known as the deuterium order parameter, S_CD_, which measures the orientation and ordering of the membrane lipid tails with respect to the bilayer normal. The order parameters at different temperatures show the ordering of the lipid hydrocarbon tails. It is computed as the ensemble and time average of the second order Legendre polynomial:

The order parameters at different temperatures show the ordering of the lipid hydrocarbon tails. Scd is measured in deuterium NMR experiments as:

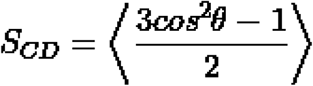

where, θ represents the instantaneous angle between the CD bond and the bilayer normal. The angular brackets denote time and ensemble average. In simulation, θ is measured as the angle between the CH bond and the bilayer normal. The more the membrane is fluid-like, the lesser the tail ordering. The S_CD_ values of the two lipid tails (averaged over all the lipids in the system and over the production trajectory) are shown in Figure 5a as a function of backbone carbons. The average S_CD_ values of the two hydrocarbon tails (averaged over all the tail atoms) are shown in Figure 5b as a function of temperature. As expected, the S_CD_ values decreased with temperature indicating increased disorderedness of the lipid tails. At 10 °C the highest S_CD_ value was found to be around 0.4 which decreased with increasing temperature becoming negligible at 50 °C. The transition temperature of DPPC and DPPG phospholipids from gel to a liquid crystalline state (*T*_*c*_) has been found to be 41 °C [57]. The sharp decrease in the S_CD_ values beyond 40 °C clearly captures this transition in our simulation. This also indicates that the presence of a single M protein dimer does not affect the intrinsic melting characteristics of the membrane lipids. However, to accurately assess the effect of multiple M protein dimers and precisely determine the transition temperature of these lipids in presence of the M protein, one has to perform atomistic simulation of model membrane with multiple M protein dimers and sample the temperature space in closer intervals, which is beyond the scope of the current work.

**Figure 5.**
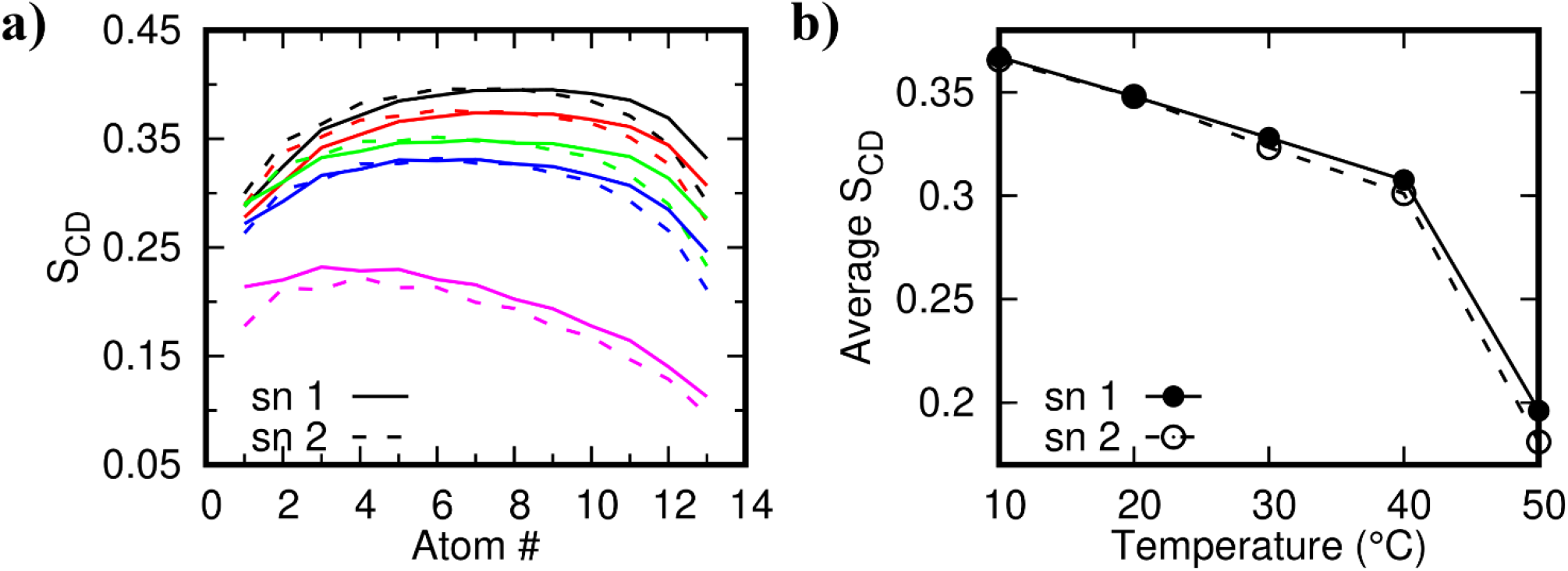
a) Deuterium Order parameters (SCD) of the two lipid tails at different temperatures displaying the relative loss of ordering at high temperatures. b) Average SCD value as a function of temperature.

While S_CD_ values indicate the ordering of the individual lipids, the relative ordering of the lipids in the bilayer can be captured in terms of the area per lipid (APL) and bilayer thickness. Figure 6 indicates the effect of temperature on these two quantities. It was found that the APL increases with temperature, while the thickness decreases. Similar to the trend observed in S_CD_, the change in APL and thickness is most prominent in the 40 - 50 °C window, again indicating the melting transition of the lipids. The experimentally reported APL value for pure DPPC bilayer at 50 °C is 0.64 nm^2^ [59–61], while that of DPPG at 50 °C is 0.67 nm^2^ [62, 63]. Thus, as expected, for a mixed DPPC/DPPG membrane with a transmembrane M protein dimer, the calculated APL values in our simulations are systematically larger than a pure DPPC bilayer [64]. Bilayer thickness of pure DPPG membrane at 50 °C is reported to be 3.55 nm [63]. It is reported to be 3.43 nm for pure DPPC bilayer at 42 °C [65] and 4 nm at 47 °C [66]. Similar to APL, the observed thickness values in our simulation are systematically larger than the corresponding pure component lipid systems. The increase in thickness can be attributed to the presence of the transmembrane domain of the M protein dimers which, owing to the hydrophobic mismatch between the bilayer and the transmembrane region of the protein dimer, can locally increase the membrane thickness [66, 67]. Additionally, a larger bilayer thickness, in general, indicates more ordered lipids and therefore, there is a possibility that the M protein dimers induce ordering in the neighbourhood lipids. However, we have not explored this aspect of protein-membrane interaction in this study and plan to do that in a comparatively larger membrane.

**Figure 6.**
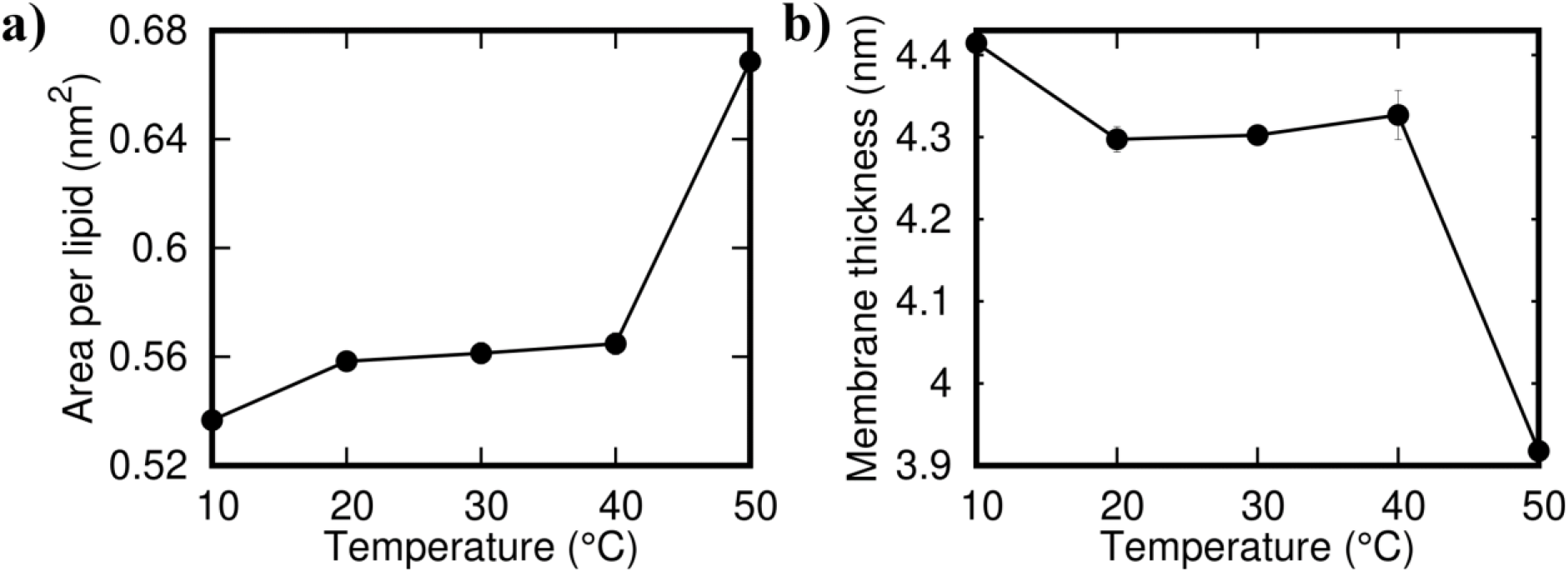
a) Area Per Lipid and b) local membrane thickness at different temperatures

Next, we examine the ability of the M protein dimer in generating membrane deformation. The mean curvature profile of the atomistic membrane at 10, 30, and 50 °C are shown in Figure 7 top panel. Here we focus on the membrane leaflet that contains the N-terminal region of the M-dimer and the mean curvature is calculated w.r.t. the membrane mid-plane, i.e., a positive mean curvature indicates the membrane to bend outward, while a negative mean curvature indicates the membrane to bend inwards w.r.t. the mid-plane. The middle and the bottom panel show top and side view of representative membrane configuration along with the position of the protein dimer. The bilayer is represented as grid points based on which the curvature and thickness calculations are performed, where the grids correspond to the lipid head groups. The bilayer grids are colored based on the membrane thickness, where the color red represents lower membrane thickness and the color blue represents higher thickness values. The mean curvature and thickness profiles are calculated over the last 10ns trajectory using g_lomepro. In the g_lomepro analysis, the curvature values are scaled by a factor of 100 for better representation and a high q-filter value of 1 is used. We found the extrinsic N-terminal domain to impart a high negative mean curvature to the membrane, thus bending the membrane inwards. A similar observation of M protein induced local membrane deformation has been made in another recent study [20].

**Figure 7.**
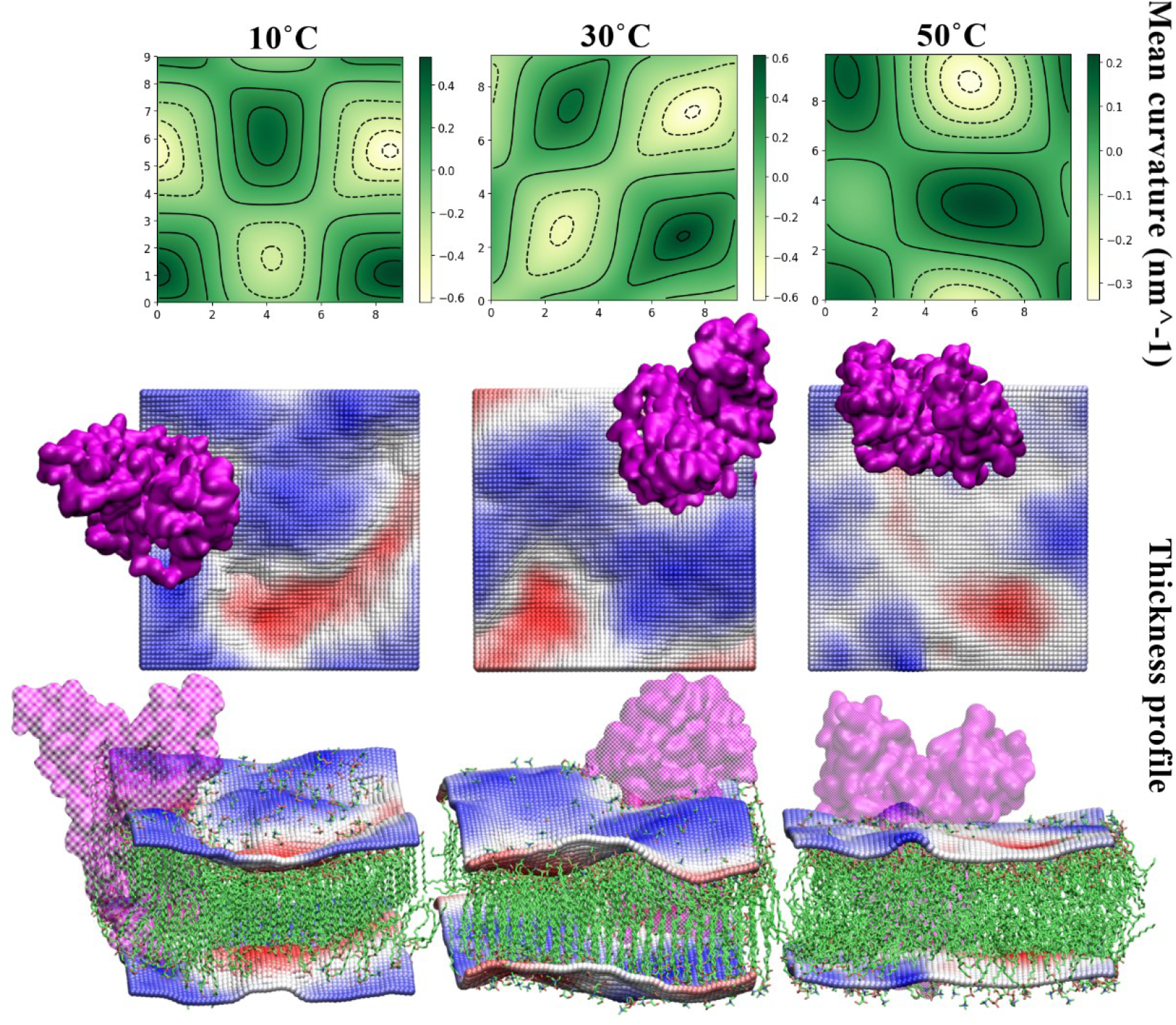
Comparison of membrane curvature and thickness at 10, 30 and 50 °C. Snapshots at corresponding temperatures are shown in the bottom two rows, with membrane color coded based on the relative bilayer thickness.

The magnitude of this bending is high at 10 and 30 °C, but almost half of that at 50 °C, which can be attributed to the increased disorderedness of the lipids and thermal fluctuations. Along with the high negative curvature at the dimer position, we also found other regions with negative and positive curvatures. The thick transmembrane region of the protein dimer comprises long helical domains that are bent w.r.t. the membrane normal (see Fig 1). Such a structure, owing to the hydrophobic mismatch, can locally perturb the lipid ordering and generate curvature. A closer look at the membrane snapshot indicates locally disordered small lipid domains even at 10 and 30 °C, which in the absence of the protein is expected to be in a gel-like ordered state. Around these small regions, the membrane exhibits lower thickness and thus, seems bent inwards (negative mean curvature). The regions with ordered lipids exhibit high thickness and thus a positive mean curvature. Due to large thermal fluctuations, the lipid membrane is fully disordered at 50 °C and the variation in mean curvature and thickness profile are less drastic as compared to the low temperature case. The extent of membrane deformation in our study is comparable to that reported in another recent study [20]. However, we must add that our system size is rather small to rule out any finite size effects and therefore, we analyzed the mean curvature in presence of multiple M dimers in CG simulation (discussed later).

### Effect of temperature on protein aggregation

Considering the pleomorphic nature of the Coronaviruses, the conformation of the M protein and the dynamics of the membrane lipids at different temperatures might impact the shape of the virions formed. Previous studies have reported that under confinement, the M proteins are more prone to aggregation [68]. To understand if temperature influences M protein aggregation, CG MD simulations were performed using MARTINI 22 on a larger system consisting of 8 membrane protein dimers (16 monomers) embedded on a lipid membrane (see materials and methods for technical details). The final conformation of the proteins at each temperature are shown in Figure At 10 and 20 °C, we did not observe any aggregates within the simulation time scale. This could be attributed to the high degree of ordering of the membrane lipids as shown in Figure 8b. At 30 °C formation of M-dimer clusters was observed within 100 ns time period, further aggregation took place within 300 ns beyond which, no more protein clusters could be observed. At 40 °C, the proteins started forming clusters within the initial few nanoseconds after which steady aggregations took place at 200 ns and 250 ns respectively. Barring two dimers, all the remaining protein dimers aggregated to form a single cluster comprising 12 proteins. At 50 °C a very fast aggregation pattern was observed and two separate clusters could be noticed.

**Figure 8.**
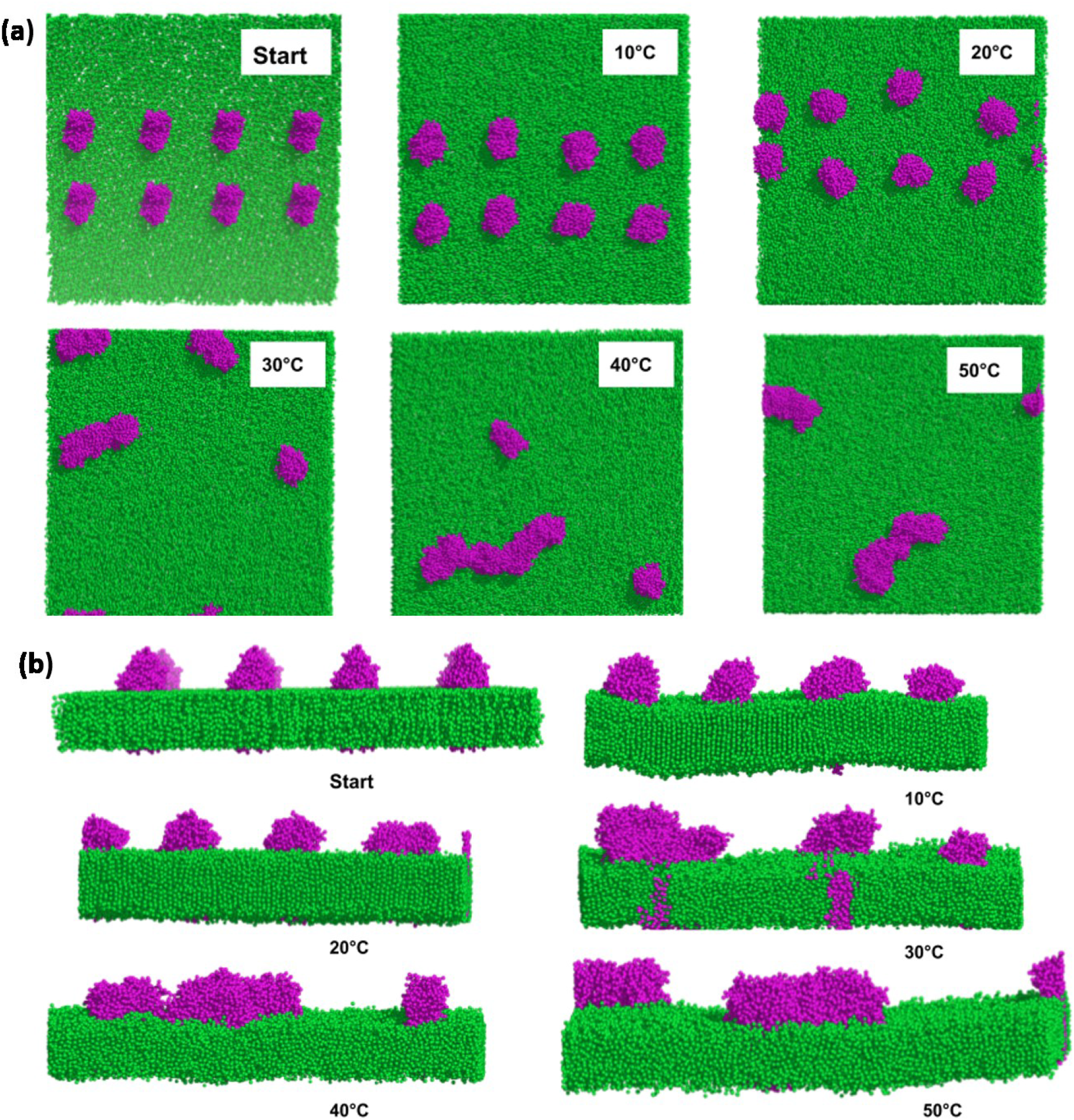
CG simulations of eight M-protein dimers (in magenta) embedded in a lipid membrane (in green) at different temperatures displaying the clustering pattern of the proteins at various temperature (a) top view, (b) side view. Note the relative ordering in lipids in (b).

The rate of formation of aggregates directly depends on the ordering of the membrane, which allows migration of protein molecules. At 50 °C due to the large disordering of membrane lipids, the aggregates were able to form very quickly (Table 1) (Figure 9). This also implies that irrespective of the noticeable conformational flexibility of the M protein at 10 and 20 °C (Fig 3), the formation of protein clusters is largely governed by the ordering of the membrane. In conclusion, we observe that clustering of M protein dimers readily occurs only beyond 30 °C.

**Figure 9.**
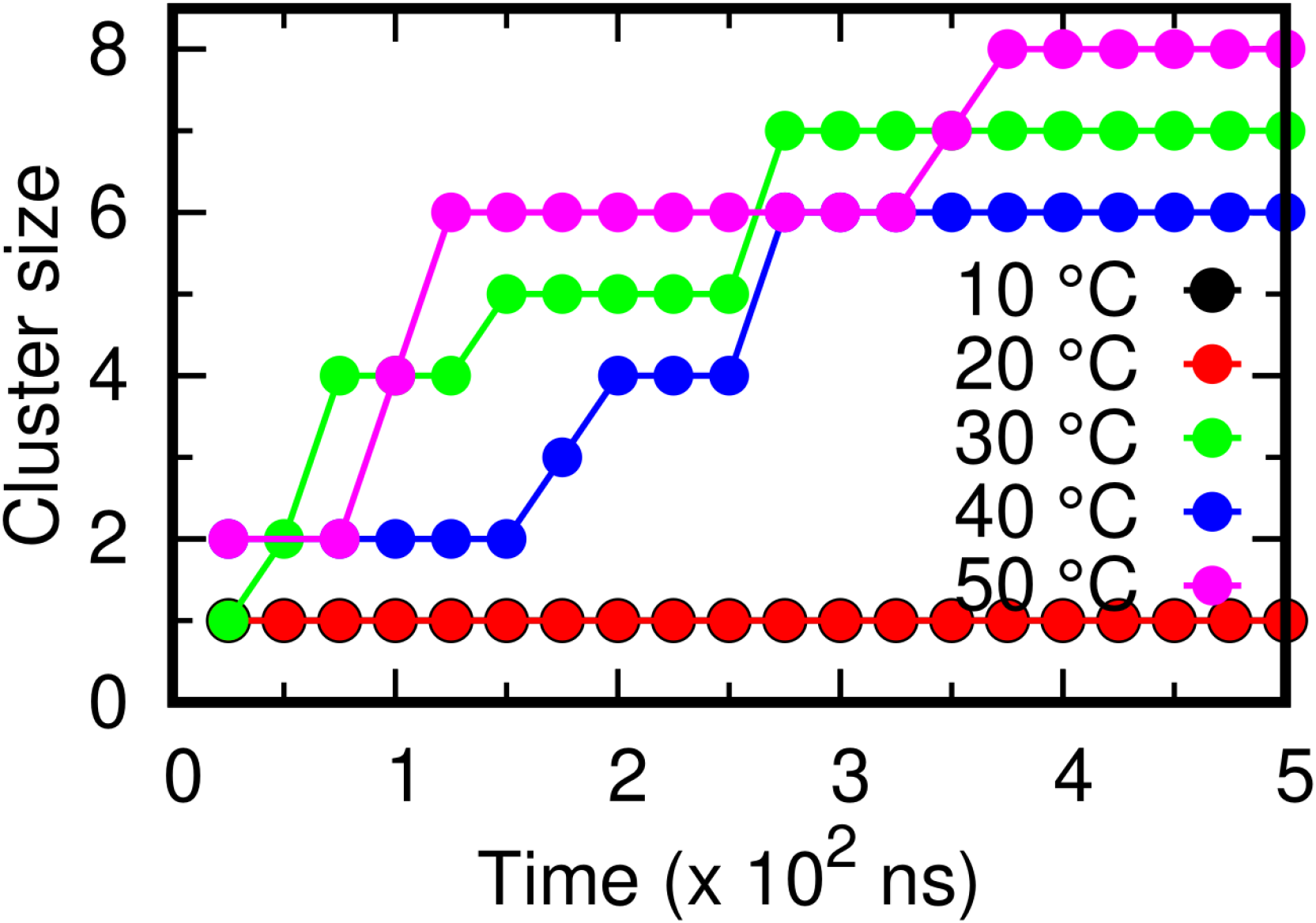
The total number of protein clusters formed at different temperatures during the 500 ns CG simulations. Colour coding similar to Figure 2.

To further clarify our observations on clustering of M protein dimers and the resulting membrane deformation, we carried out CG simulation involving 64 dimers on a lipid bilayer. We specifically focused on the clustering behaviour at 30 and 40 °C and the resulting membrane curvature. We show the final membrane conformation and the resulting mean curvature profile in Figure 10.

**Figure 10.**
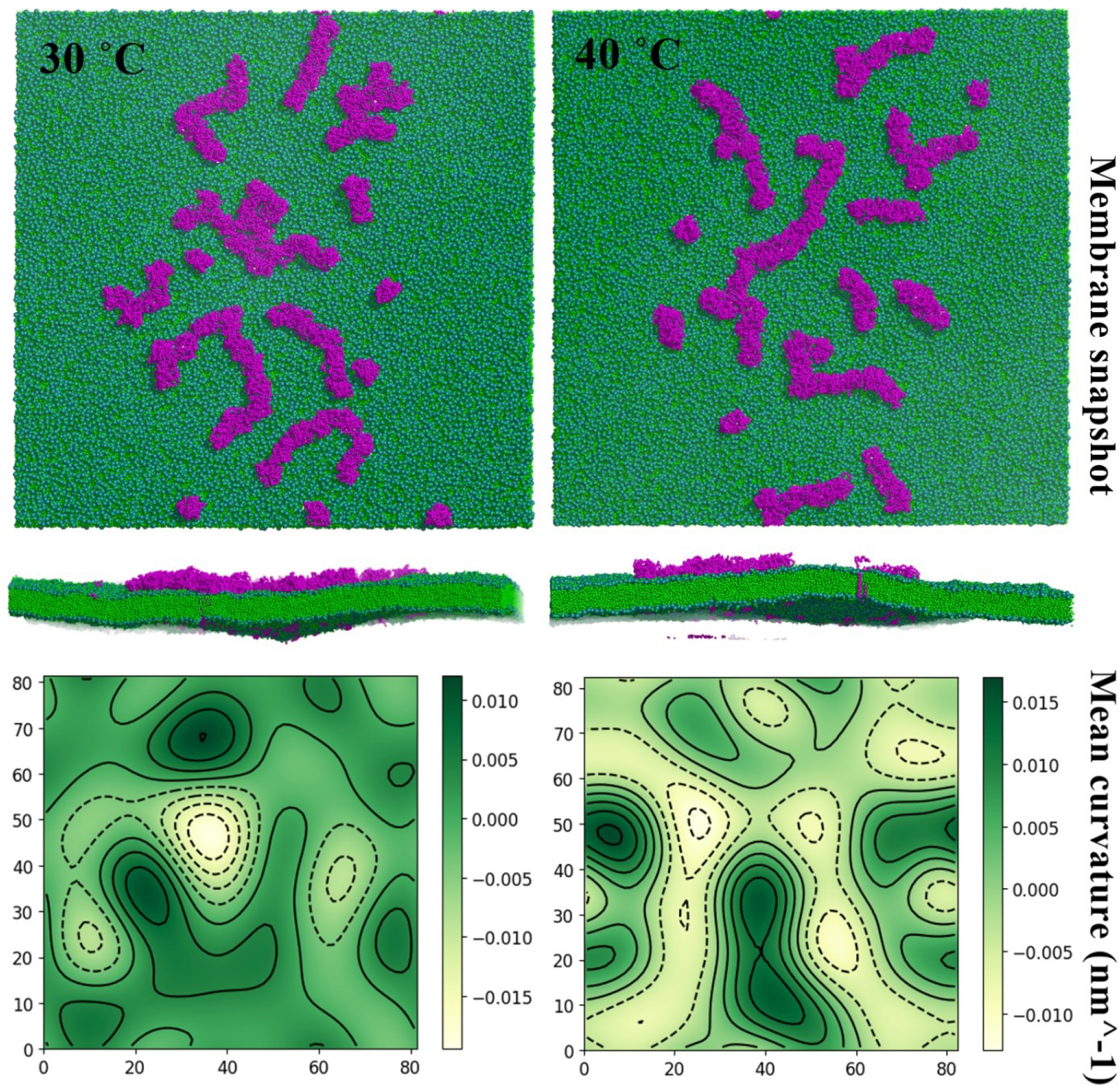
Comparison of membrane curvatures (bottom panel) with 128 M protein monomers at 30°C and 40°C. The top panels show the top and side views of a representative membrane snapshot at each temperature.

While we could not observe all the dimers clustering to form a single large cluster during the time scales of the simulation, the effect of the partially formed clusters on the membrane curvature was evident. Though the presence of these clusters induced appreciable local curvature, we did not observe any large-scale membrane deformation at either temperature. This observation suggests that M protein alone may not be capable of generating large curvatures that can trigger budding/fission events and other integral proteins might play a crucial role. However, owing to their aggregation capability, the M protein might facilitate the curvature generating process. We will focus on investigating this collective mechanism in our future work.

## Conclusion

Out of the four different structural proteins of the SARS-CoV-2 virus, the M protein is the most abundant and widely distributed. It associates with the S, E and N protein and also with other M proteins and directs the virus assembly and aids in the viral release. The pleomorphic nature of the coronaviruses allows them to change the particle size and shape, which also depend on the structural proteins of the SARS-CoV-2, especially the M protein. Hence, understanding the dynamics and behaviour of the M protein is very important to understand the formation of the virus particle. Although the variations have been reported in literature, very few studies have been carried out to understand how the overall virus structure is shaped.

We performed atomistic and coarse-grained (CG) simulation on ionic DPPC/DPPG lipid membranes embedding M protein dimers to understand the effect of temperature on the behaviour of both the protein and the membrane, and their mutual interactions. The atomistic simulation on single M protein dimers indicated that the conformation and dynamics of the M protein depends strongly on temperature, as expected. As both DPPC and DPPG lipids exhibit a transition temperature of 41 °C, the transmembrane domain was found to be flexible between 30 and 50 °C, where the transmembrane and extrinsic domain of the proteins could sample various conformations. PCA performed on the extrinsic N-terminal domain of the protein dimer further indicated that the free energy landscape of the dimer goes through significant change over this temperature range. The energy landscape at 40 °C was found to exhibit some well-defined basins indicating the possibility of functionally important dynamics that occurs over the physiological temperature range. While the structure and ordered/disorderedness of lipids seem to play a crucial role in dictating the structure and dynamics of the embedded protein dimer, the overall nature of the lipid membrane was not much affected by the presence of the protein. The order parameter, APL and bilayer thickness, all showed signatures of the transition temperature beyond 40 °C. However, we could observe the overall bilayer thickness to be affected by the protein, which can be attributed to the hydrophobic mismatch between the lipid bilayer and the protein transmembrane region. We also observed some interesting interplay of protein-membrane interaction in the mean curvature profile of the membrane. While the protein locally induced negative mean curvature, it also induced appreciable positive and negative curvature around it. With increasing temperature, these fluctuations die down. Our observations, thus, suggest that the protein might lead to ordering/disordering of the neighbourhood lipids, which further induces local curvature. However, to characterize such ordering, a systematic analysis of the lipid dynamics has to be performed in order to identify local ordered/disordered regions and their correlation with the protein. Finally, our system might be small to clearly capture any large-scale undulations and finite size effects might also have some effects.

To better understand the effect of temperature on protein clustering and the resulting membrane curvature, we performed CG simulations. Simulations on 8 protein-dimers indicated that protein clustering happens only at 30 °C and beyond and the rate of clustering increases with increasing temperature. This again should be attributed to the gel-like ordered phase of the lipid membrane. Simulations on 64 protein dimers at 30 and 40 °C further indicated the effect of protein clustering on membrane curvature. While the dimers failed to form a single large cluster during the simulation time scales, the partially formed clusters could induce appreciable membrane curvature. However, the scale of the deformation was far from that expected in a case where one can expect budding. As M proteins of SARS virus are implicated in virus budding, it remains unclear if M protein alone is capable of leading to such large-scale deformation. In our study, we find this to be not feasible.

As we observe the most interesting changes in the protein-membrane system happening close to the physiological temperature range, we believe that the clustering of M protein dimers and the resulting change in membrane curvature might be an important factor in the viral activity. However, as the viral membrane consists of various other proteins as well, the behaviour of M protein dimers in presence of other proteins can provide crucial information on their mutual role in the viral life cycle. Recently, such a study has been performed [20], which aims at understanding the role of M and E proteins in generating membrane curvature. They found the M proteins to cooperatively generate membrane curvature, while E protein kept the membrane relatively flat. An important aspect in the membrane-protein study is the model membrane which should closely represent the viral membrane. In this line, the study by Monje-Galvan and Voth [18] shows the difference in the lipid sorting pattern and the membrane deformation profile around different structures of the M protein. It will be interesting to extend our study to understand the effect of temperature on cooperative protein-protein and protein-membrane interactions on complex model membranes. Additionally, CG MD simulation can enable the study of large-scale protein-membrane systems, which can provide crucial information on the aggregation behaviour of the proteins and their effect on the lipid membrane.

## Acknowledgements

SLR thanks NIT Warangal for research seed grant (P1131) and National Energy Research Scientific Computing Center of the Ernest Orlando Lawrence Berkeley National Laboratory, a DOE Office of Science User Facility supported by the Office of Science of the U.S. Department of Energy under Contract No. DE-AC02-05CH11231 and the Extreme Science and Engineering Discovery Environment (XSEDE). We are grateful to the Covid19 HPC Consortium for providing resources and helping researchers work for a noble cause. We would like to thank Dr. Shikha Prakash and Prof. Durba Sengupta for their useful suggestions.

## Author Contribution

SLR designed research; SLR and MT performed research; SLR, MT and NM analysed data; SLR, MT and NM wrote the paper.

## Additional Information

### Supporting Information Available

Includes Table 1 and Figure S2. This information is available free of charge via the internet

### Competing Financial Interests

The authors declare no competing financial interests.

## Notes

### Competing Interest Statement

The authors have declared no competing interest.

