## Supplementary material for "How does temperature affect the dynamics of SARS-CoV-2 M proteins? Insights from Molecular Dynamics Simulations": Supplemetary Information

**Table S1.** Details of Systems

| Serial No. | System | No. of M protein | Simulation Time |
|---|---|---|---|
| 1 | All atom | 2 | 500ns |
| 2 | All atom | 2 | 500ns |
| 3 | All atom | 2 | 500ns |
| 4 | All atom | 2 | 500ns |
| 5 | All atom | 2 | 500ns |
| 6 | Coarse Grained | 16 | 500ns |
| 7 | Coarse Grained | 16 | 500ns |
| 8 | Coarse Grained | 16 | 500ns |
| 9 | Coarse Grained | 16 | 500ns |
| 10 | Coarse Grained | 16 | 500ns |
| 11 | Coarse Grained | 128 | 200ns |
| 12 | Coarse Grained | 128 | 200ns |

**Table S2. Total number of atoms**

| Serial No. | System | No. Of M Proteins | No. Of DPPC | No. Of DPPG | No. Of Solvent |
|---|---|---|---|---|---|
| 1 | All atom | 2 | 210 | 90 | 29273 |
| 2 | Coarse Grained | 8 | 3423 | 1467 | 112172 |
| 3 | Coarse Grained | 16 | 14791 | 6339 | 493141 |

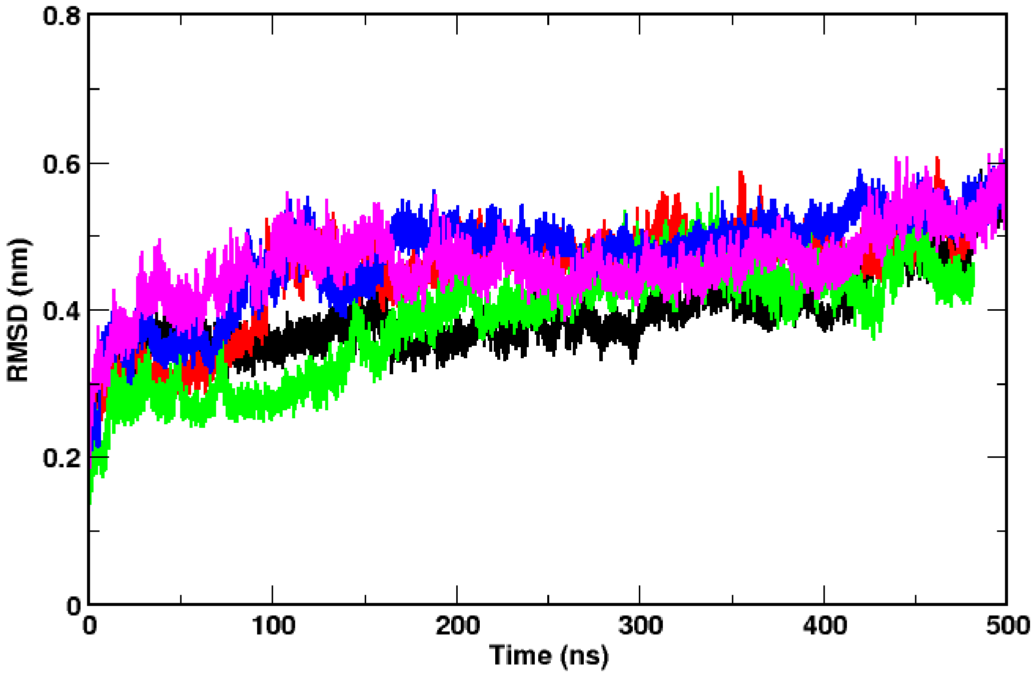


**Figure S1.** RMSD of M protein dimer in atomistic simulations at 10˚C, 20 ˚C, 30 ˚C, 40 ˚C and 50 ˚C
